# Native DNA electronics: the Nuclear Aggregates of Polyamines possible role

**DOI:** 10.1101/294199

**Authors:** L. D’Agostino

## Abstract

The genomic DNA is enveloped by nanotubes formed by the Nuclear Aggregates of Polyamines (NAPs) that induce DNA conformational changes, and provides protection and increased interactive abilities for the double strands. In a physiologic environment, the nanotube arrangement initiates with the spontaneous interaction among the terminal amino groups of polyamines and the phosphate ions, with the consequent formation of cyclic monomers that hook at DNA grooves. The polymer thus formed has the morphological features of an organic semiconductor and, therefore, can be considered able to conduce electric charges. Phosphate ions positioned on NAP external surface could regulate, as in a physical electric circuit, both protein linear and rotational (histones) motion, in accordance with the basilar principles of the electronics. A model of a carrier system for proteins motion along the polymer wrapping the DNA strands, based on the phosphate-phosphate complexation, is proposed.

## INTRODUCTION

In recent times, DNA has been considered an interesting model of an organic circuit, limitedly to the bases compartment of its structure. The rings of purines and pyrimidines, heterocyclic aromatic compounds, are apt to conduce electrons and, therefore, have been proposed as semiconductors. The presence of electronic delocalization that characterizes π-π moieties is fundamental for the transfer of the electrons, but the low or high conductivity of the π-stacked polymers depends on the ordered alignment of aromatic moieties since their semiconducting features are originated from intra-supramolecular interactions (1,2).

Not all base couplings are efficient in charge transfer since a significant semi-conducing activity was limited to the guanine-guanine one. In particular, the best conductivity was reached by the stacking of alternative series of five guanines. However, the further elongation of guanine blocks was not able to guarantee a coherent charge transport across the DNA (3).

All these factors indicate that native DNA bases are not ideally connected in order to assure an efficient charge transfer, as necessary for a true circuit that has to work physiologically. Another factor that precludes the charge transferring is the alignment of the moieties onto the double strands that are, anyway, essentially "insulated", being the periphery of the bases flanked by the poor conductor sugars (4).

All the studies concerning the DNA bases conductance confirm that the stacking of π-moieties is fundamental for the development of DNA nanotechnology devices (5), but also evidence that a long-range charge transport cannot happen in naked unmodified double strands. Therefore, a role of electronics in DNA physiology cannot be acknowledged, if one limits its consideration to the setting of the bases. On the other hand, another "aromatic" environment is present onto the strands: in native conditions, they are enveloped by a polymeric system of nanotubes formed by the self-assembly of polyamines and phosphate in cyclic structures the Nuclear Aggregates of Polyamines (NAPs) (6-15) - that I suggest here as a powerful electronic apparatus that assists the DNA in its functions.

This paper has the aim of interpreting the ascertained morphological and functional characteristics of NAPs at the light of the basic principles of the electronics, and hence to delineate their possible role as electronic structures acting in still debated fields of cell nucleus physiology.

## METHODS

The electronic functioning of NAPs was theorized by the insertion of the biochemical, biophysical, analytical and structural produced data in the setting of the current bio-electronic knowledge, at light of the basic principles of electronics and mechanics. The ideation process has been expressed in drawings by means of AutoCAD by Autodesk Inc., San Rafael, U.S.A. (Figure 1) and Windows10 graphical tools (Figure 5), respectively.

**Figure 1.**
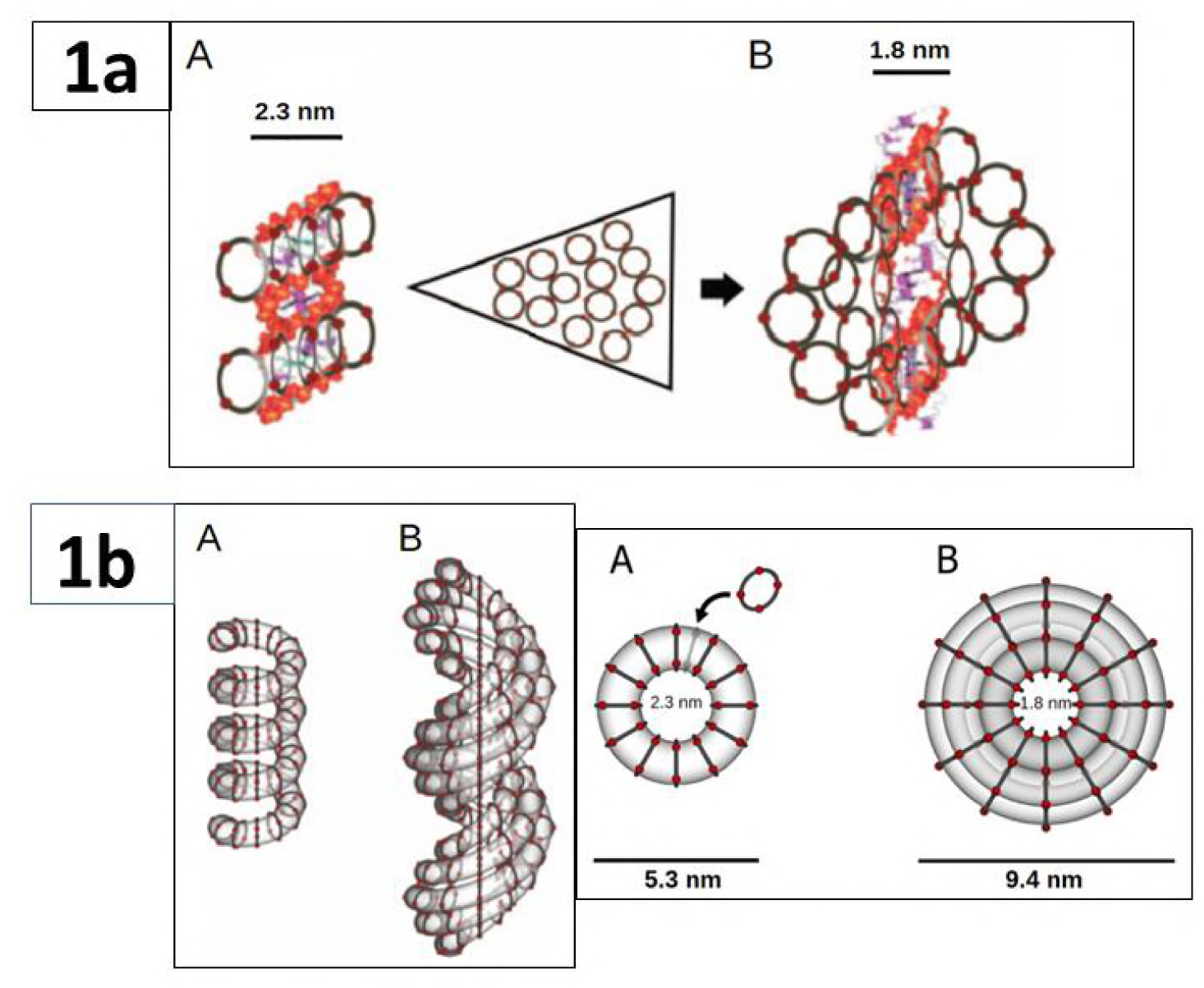
Interaction of single NAP monomers with different DNA forms. (A) s-NAP interacting with A-DNA. A-DNA has a groove width more suitable than other DNA forms to interact with this NAP. The addition of s-NAP units to two s-NAPs already bound to DNA (up to five) allows the formation of m-NAP directly onto the DNA. This may favor the transition to the Z-DNA, through the progressive widening of DNA strands and the exposure of bases. (B) The final effect is the formation of a m-NAP interacting with Z-DNA. Z-DNA form stabilization by the m-NAP arch-like structure was represented as due to the distancing of consecutive A-DNA major grooves.This model das been imagined on the basis of electrophoretic genomic DNA behavior at 37 C° and established double-strands measures and shapes (7). Nanotube models. (A) Adjacent s-NAPs are imagined to produce a tubular structure enveloping the A-DNA. (B) Adjacent m-NAPs are imagined to produce a nanotube enveloping the Z-DNA. In the bird’s eye view (right panel), the NAP monomers depicted are those connected to the phosphates of the DNA (12 per helix turn). Other NAP monomers can be imagined in parallel along the transparent sections of the tubes. The nanotube is formed by the assembly of cyclic monomers onto 12 hanging points per DNA helical turn (one for each phosphate pair per helix), and completed with accessory blocks flanking the DNA anchored rings, which are linked through additional lateral H-bonds established among the available phosphate groups and other stacking interactions. The 1b right panel is a new graphical integration of previously published images (8). The AFM analyses confirmed the models (Figure 2)

**Figure 2.**
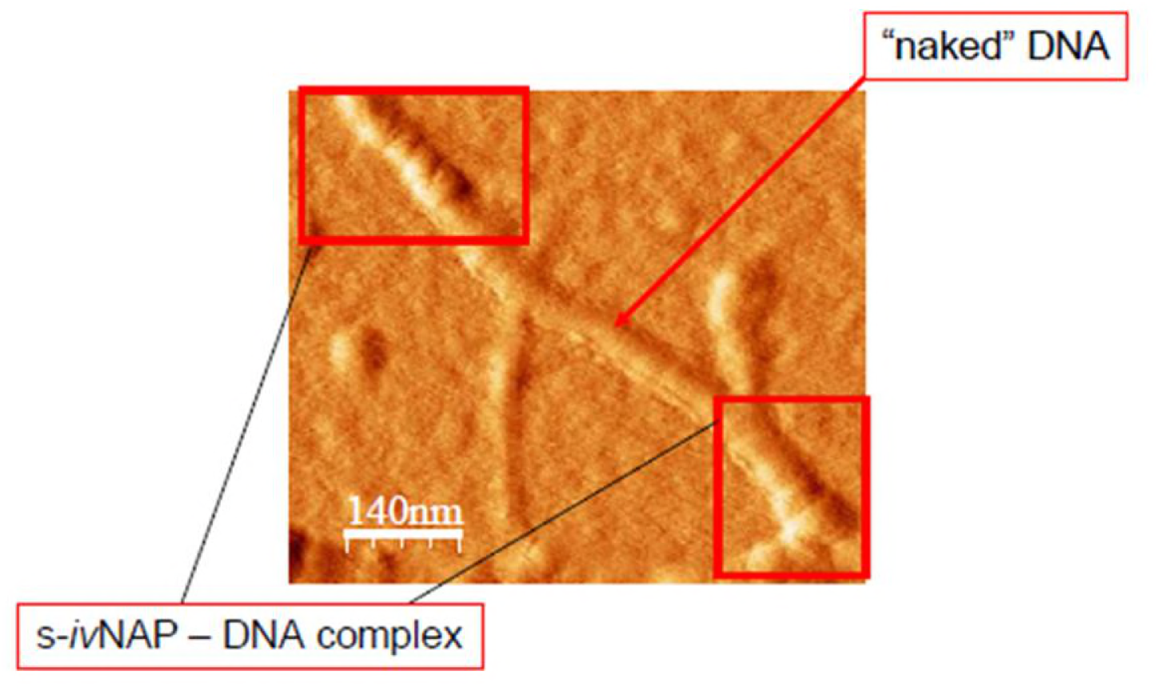
AFM image of s-NAP enveloping a double-stranded DNA. The red boxes delimit the areas covered by the nanotubes, formed by s-NAP monomers, that exalts the helicoidal motif of the DNA strands. In the central tract, the DNA is naked and some s-NAP debris, due to the nanotube damage produced, probably, by the cantilever passage, can be perceived (11). and evidenced that NAPs also mediate the interaction of the double strands with HMGB1, a DNA binding protein (10,11,13-15) (Figure3).

## RESULTS and DISCUSSION

### Nuclear Aggregates of Polyamines

Polyamines (putrescine, spermidine, and spermine) are linear polications that, in the presence of phosphates, intrinsically form hierarchical assemblies of complex structures. Polyamine – phosphate interaction, we described in 2002 (6), was successively found in other settings (16- 18).

Three compounds referred to as NAPs were isolated from nuclear extracts of many different cells (6). Polyamines and phosphate ions also self-structure in vitro within well-defined ratios under the conditions of thermodynamic equilibrium and independent of the presence of the DNA template to form in vitro NAPs (*iv*NAPs) (9). According to gel permeation chromatography, their estimated molecular masses are about 8000, 5000, and 1000 Da and, consistent with the nomenclature of extractive NAPs (7) these compounds were named l-, m-, and s-*iv*NAP (large, medium and small in vitro NAPs), respectively (9). The *iv*NAPs share structural and functional traits with their cellular counterparts (9).The polyamine-phosphates aggregates have positive net charges at physiological pH and interact with the negatively charged phosphates of DNA. As a consequence of this interaction, NAPs influence DNA conformation and protect DNA from nuclease degradation. Our results led to the conclusion that this class of aggregates fully preserves the integrity of DNA without affecting its elasticity (6-8).

NAP modeling that combined experimental evidence (6-8) with a number of theoretical assumptions (19, 20) was performed following a step-by-step ideation process. First, the three polyamines, protonated at physiological pH, ionically binding a phosphate anion at their cationic edges, cooperate to generate ring-like aggregates, as suggested by the appearance of an absorption band at λ≈280 nm that is indicative of an electronic delocalization similar to that of aromatic compounds. No absorbance at 280 nm was detected when polyamines were individually incubated under the same conditions (6,9). Second, in accordance with the principles of the hierarchical aggregation, NAPs interact with DNA grooves by arranging their rings into structures that we supposed to be similar to nanotubes that completely wrapped the double helix (7,8) (Figure 1).

All these data are in accordance with the belief that these self-aggregated supramolecular structures act as polymeric smart nanotubes that play crucial subsidiary functions, like conformation, protection, and interaction towards the double strands. Since these aggregates, firstly isolated by us in the cell nuclear, extracts (2), are strictly dependent on both a physiological pH and osmolarity, it is sure that they operate in the nuclear setting of living cells (6-8).

The circular monomers formed by the electrostatic interaction of polyamine N termini and phosphate ions, as well as their interaction with, or their self-aggregation onto, the backbone phosphates are driven by the electrostatic forces. However, H-bonds, stacking and other weak interactions regulate the alignment of the cyclic elements onto minor DNA grooves, thus forming tubular structures enveloping the entire DNA (7,8).

### Proposition of NAPs as an electronic circuit wrapping the DNA

In a semiconducting material, such as silicon, an inorganic polymer, the outer electrons of the atoms can be replaced, albeit not as easily as in a metallic conductor. Silicon is a typical semiconducting material that, with the supplementation of alien atoms (doping) assume the ability to produce additional free electrons that can be displaced to neighboring atoms, thus transferring the charge. A semiconducting material with an excess of electrons, typically obtained by phosphorus contamination, is named an n-type semiconductor. There is also a p-type semiconductor, doped with atoms that lack electrons (i.e. boron), which create electron holes in the material. On applying a voltage to a p-type semiconductor, the electron holes move from the positive pole to the negative, and a net current flows in the opposite direction. If an electric field is applied across the n-type semiconductor, electrons move from the negative to the positive battery terminal (21).

In organic semiconductors formed by aromatic moieties (22), the delocalized electrons are, in accordance with the “frontier molecular orbital theory” (23), shifted among interacting molecular orbitals in the way that the highest occupied molecular orbital (HOMO) produces the entire negative charge coming from the electron pairs and, correspondingly, the lowest unoccupied molecular orbital (LUMO) operates as an acceptor (24), so assuring a directional electronic flow, provided that a nonzero energy gap between HOMO and LUMO is assured (25-27).

Consequently, an organic circuit has to have superimposed connected monomers. In this regard, NAPs show stacking abilities also without the DNA strands support (10: suppl_file <u>bm101478j_si_001</u>). However, in the cell nucleus NAP polymeric assembly is strictly dependent on strand interaction so that mono-cyclic or pentacyclic structures can be formed in order to produce A, B, or Z forms (Figure 1a). The supramolecular organization bases on cyclic modules that have 12 hanging points per DNA helical turn (one for each phosphate pair per helix). Furthermore, since links among adjacent anchored monomers cannot be directly established, as the distance between two closest NAPs that have connection with the phosphates of the DNA is always greater than 0.3 nm, the limit beyond which the hydrogen bond (H bonds) cannot be established, it was theorized that these NAPs rings were sided by others that maintain continuity by means of H bonds among the external phosphate groups and other weak interactions.

The final effect of this assembly is a nanotubular frame enveloping the entire DNA (Figure 1b) that follows the electronic criteria of a long n-type organic semiconductor.

#### NAP functional outcome

NAPs, by means of their interaction with DNA groves, protect the strands by DNAse degradation and potentially by ɣ-radiation and provide them with dynamic assistance during their conformational changes (6-8,12).

But another important function can be ascribed to these super-aggregates: their interaction with proteins. (15). AFM images clearly show that HMGB1 single molecules assume a pearl necklace configuration, being aligned onto the external wall of the nanotube (Figure 3).

**Figure 3.**
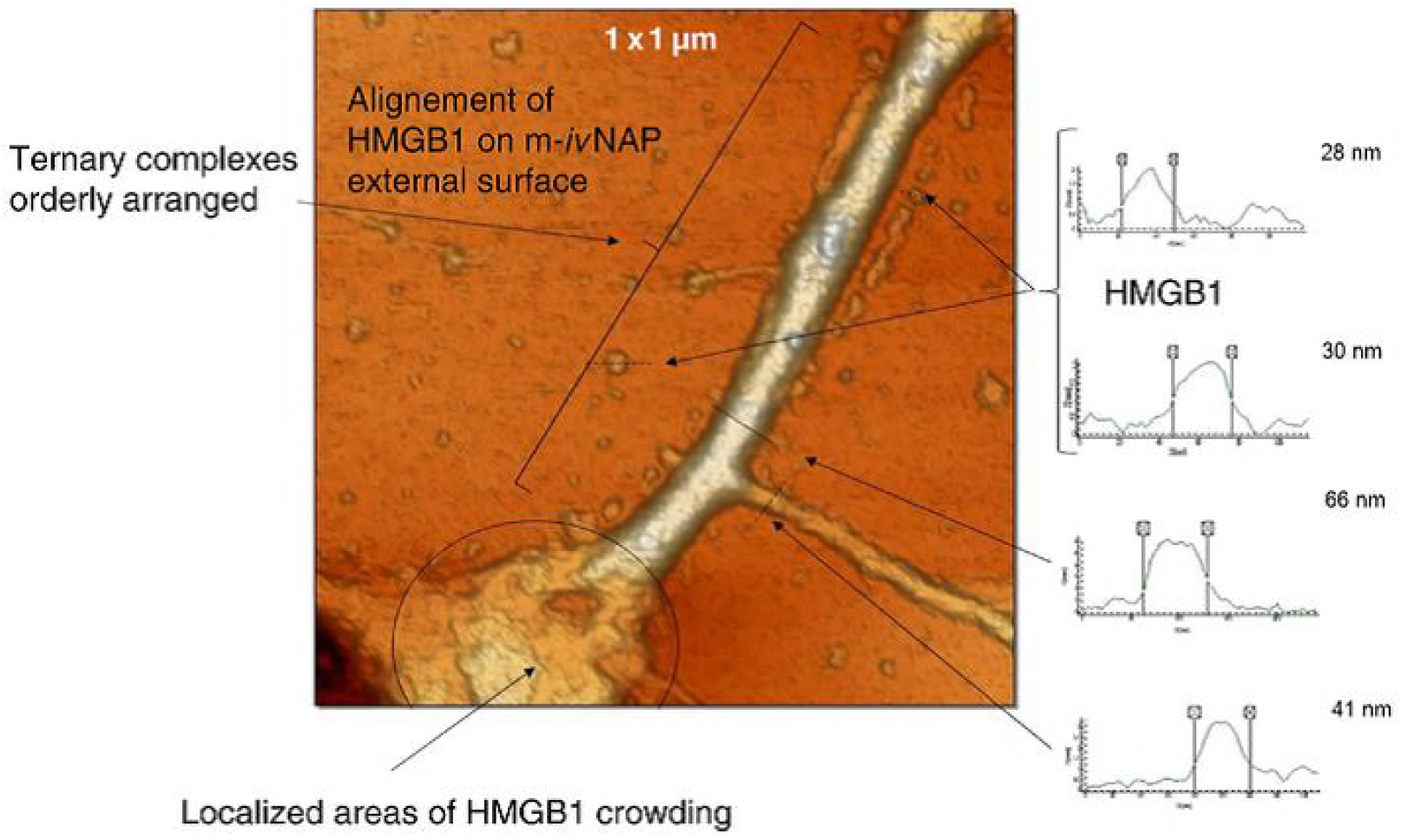
AFM image of a Z-DNA tract covered by the m-NAP nanotube, further complexed with HMGB1 proteins (Ternary complexes). These proteins self-dispose in a pearl-lace motif at a few nanometer distance from the strand, so indicating that this space is occupied by the m-NAP nanotube (15).

Protein-DNA super-aggregation is permitted by the interaction of the positive charges of the proteins and the external NAPs’ phosphates. In fact, the outer surface of the nanotubes has a charge asset utilizable for protein interaction, due to the presence of phosphates-phosphate complexes (13,28,29) (Figure 4).

**Figure 4.**
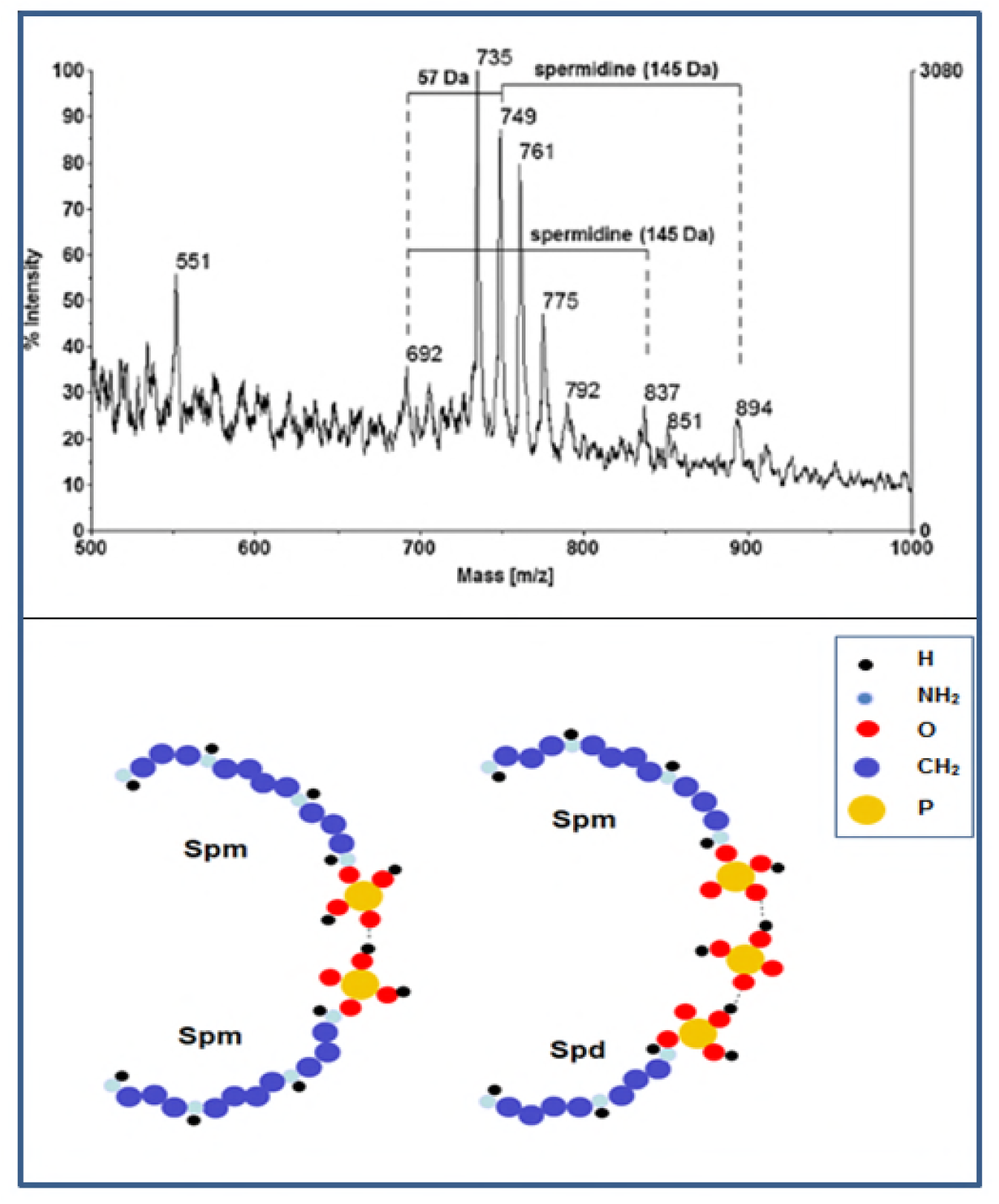
MALDI-TOF MS signals of s-ivNAP and their possible assignment (only the external face of s-NAP is represented). Spm: spermine; Spd: spermidine (13).

This ability leaves the door open to speculation about a possible implication of the phosphate-phosphate complexes locked by H bonds in protein motion along the strands that will be presently discussed.

### The role of electric charges in protein motion

Proteins motion along the DNA consents their rapid arrival on the own interactive site determined by a specific base sequence. The rush to the DNA target site is an exceptionally rapid process influenced by DNA structural parameters, like the curvature and the extent of helical twisting, that concur with the shape of the groove widths and the electrostatic potential around the DNA molecules (30). From a charge density point of view, the interaction of HMGB1 with NAPs, recently reported by us (15) is expected to be mediated by negatively charged phosphate groups located in their outer part. In accordance with Mata et al. (29,31), the external surface of NAP monomers interfaced with the DNA, exposing two or three H-bonded phosphates, guarantee the maintenance of a negative charge cluster in the setting of positive (protonated) charges (13,14).

Inter-phosphate interactions have an intrinsic stability that explains the easy aggregation of these anions in spite of ionic repulsion. In solution, in fact, solvent molecules screen ionic repulsion, effectively removing the energetic barrier that could hinder the formation of the complex (32).

The H bond is a very strong link, only surpassed in power by that produced by ionic forces. The phosphates H bonds, differently from ionic ones, are not significantly influenced by the environmental electric field (EF) depending on adjacent solute cations, so becoming dominant in the supramolecular structuring (31).

The localization and movements of a messenger protein are highly regulated by Coulomb interactions between both a radially directed EF (33) from the cell nucleus into the cell membrane and the net protein charge determined by its isoelectric point, phosphorylation state (each phosphate adds roughly 2 negative charges), and the cytosolic pH. Namely, due to the Coulomb interactions of the phosphorylated negatively charged dominions with an intracytoplasmic EF, messenger proteins were found to move rapidly (< 0.1 sec) from cell membrane to the perinuclear cytoplasm (34).

By these evidences, it seems clear that the classical physico-chemical variables involved in protein electrophoresis, rule also the mechanisms for the rapid shuttling of messenger proteins.

It is worth mentioning that the methodology used in both these above-mentioned studies (33,34) was very respectful of cell integrity: the EF was evaluated by 30 nm ‘‘photonic voltmeters’’, 1000-fold smaller than traditional voltmeters, which enabled a complete three-dimensional EF profiling throughout the entire volume of living cells. These nanodevices, which can be calibrated externally and then applied for the EF determinations inside any live cell or cellular compartment, ascertained that the EF (−3.3 × 106 V/m) from the mitochondrial membranes penetrates much deeper into the cytosol than previously estimated. In particular, the EF associated with the polarized mitochondrial membrane dropped significantly and rapidly with distance from it and, although the cytosol EF intensity never achieved the maximal intensity measured at the mitochondrial level, the EF was still measurable several microns away from the mitochondria (33). Cunningham et al. (34), too, evidenced the decline in the EF with distance from the nuclear membrane, having determined a radially directed EF from the nuclear membrane into the cell one, as well as its local perturbation caused by the presence of mitochondria due to spontaneous transient depolarizations in mitochondrial membrane potential.

Transient openings of the mitochondrial permeability transition pore (mPTP) (35,36) are believed to produce these rapid changes of the membrane potential (ranging from <10 mV to >100 mV) named mitochondrial flickers (37).

A similarly refined methodology was not specifically applied to the cell nucleus setting. Loewenstein et al., using two microelectrodes impaling the nucleus of a Drosophila salivary gland cell, (38) and Mazzanti et al., by means of an enucleation procedure in isolated murine pronuclei (39) detected negative intranuclear potential differences. These results were confirmed by Oberleithner et al. that found in nuclei of giant MDCK cells (in reference to the cytoplasm) an intranuclear potential of -4.1+0.9 mV, ascribed to the high concentration of nuclear acidic proteins and/or to negative electrical changes of the phosphate backbone of the DNA (40). However, nuclear membrane has a recognized, although not fully defined, role in conditioning the nucleus electronics. Namely, the transport of macromolecules and small ions is handled by the nuclear envelope with its nuclear pore complexes (strictly ruled supramolecular structures) by means of two separate physical pathways, the large central channel and the small peripheral channels, respectively (41).

In conclusion, it is evident that protein motion in the cytoplasm, as well as, in the nucleus, is ruled by the EF and that the Coulomb interactions among proteins and ions present in their microenvironment play crucial roles in the mechanism.

The biochemical structure and the binding forces involved in NAPs system seems to fit well with the factors regulating the messenger proteins motion in the cytosol (33,34). As can be evinced by HMGB1 behavior (15), the interaction of protein with a phosphorous dominion (the external NAP surface) and a guiding EF (assured by the NAP semiconductor device) can be the principal dynamic factors at play in the nucleus also. Theoretically, the efficiency of a possible motive apparatus should be also greater around the strands than in the cytoplasm: in fact, while for cytosol proteins the run happens in the three dimensions (34), in the nucleus, the DNA-binding proteins could be guided by the NAP phosphate-phosphate track (13-15), so having a strictly predetermined, finalized and efficient motion.

### NAPs as electronic devices for protein motion

Theoretically, the intrinsic mechanisms of protein shuttling along the external face of NAPs reside on its phosphates rich external surface. As can be seen in Figure 3, in m-*iv*NAP covered DNA, HMGB1 are aligned in the external face of the polymer, so implying an underneath row of phosphates that interact with the amino terminals of the protein.

In Figure 5, showing a model of a carrier device for binding proteins, phosphates in line are subjects to both coulombic repulsive forces and inter-phosphates H bonds. The selective property of this structural configuration arises from the high directionality of the involved H-bonding interactions that oppose rerouting to other self-assembly pathways (42).

**Figure 5.**
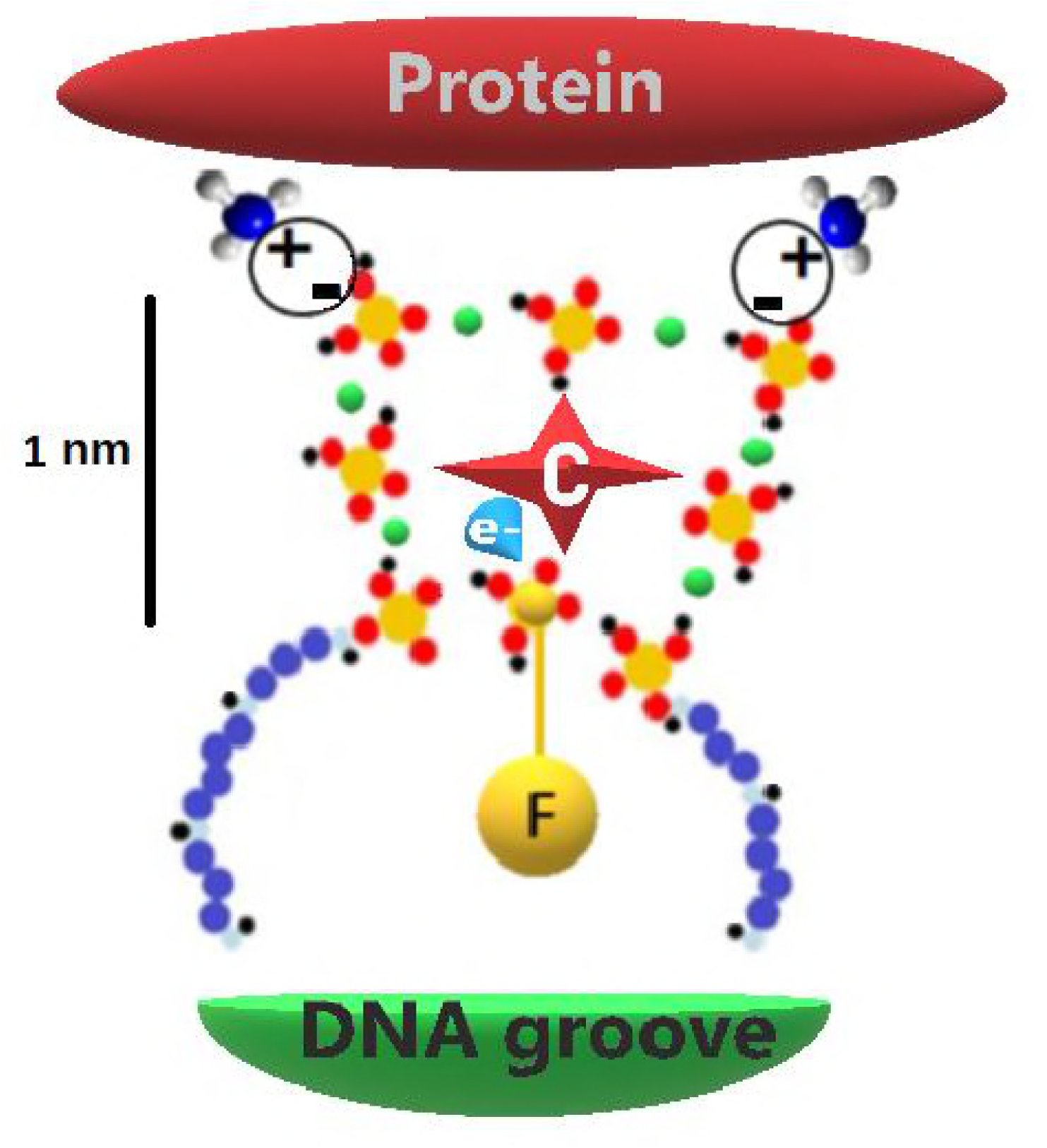
This is a design of the elemental components, and their bonds, involved in a carrying system made of phosphate-phosphate complexes located on the external surface of a NAP nanotube interacting with a protein having amino-groups available for establishing ionic bonds with the phosphates.The phosphate octamer is held by hydrogen bonds (green dot), but the central phosphate group, labeled with F, is able to stream freely in an oriented electric field (blue point of the arrow with e-). Protein motion is produced by the repulsive Columbic negative forces (C inserted in redstar) sustained by the streaming of F phosphates, whilst the H-bonded phosphates maintain the protein rush in a sort of track. Protein stops the run when its appropriate base sequence in the DNA groove is reached.(For clarity reasons: 1. only the external edge of NAP monomer is depicted; 2. the dimensions of polyamines are underestimated in order to graphically exalt the phosphates assembly; 3. therefore, the length of the mark refers only to the phosphate-phosphate carrier).

The phosphate–phosphate arrangement (Figure5), under the "directional pressure" of the electronic circulation in the NAP circuit, produces a linear flow of phosphate ions that transports the protein along the DNA up to the recipient base sequence, that exerting specific attractive forces, finally arrest its run.

#### Linear motion

The function of H bonds in this kind of motion that can be defined of the type "projectile motion in the electric field" (which tends to have a parabolic path) is to keep the shuttle in an adequate track. The allowance of the transported protein on the phosphate bogie is permitted by coulombic attractive forces and the locomotion is sustained by coulombic repulsive ones. However, even if the electrical force of this transport mechanism is based on the Coulomb law, the influence of the charge density modifications depending on the various physiological tasks and the braking and restraining role played by H bonds are influencing factors.

Phosphates occupying a central position in the phosphate-phosphate construction maintain the double negative charge and, even if ideally have the possibility of sharing single H bonds per side, have a so high degree of mobility that they constitute, in reason of the repulsive coulombic force, an unrestricted streamflow of negative charges (in Figure 5, the mobile phosphate is marked with F). Only the single elements contributing to the carrier assembly were here represented in this image. Actually, the system should be much more composite since, in the NAP induced Z-DNA form, which is the type of DNA involved in active tasks, like gene expression and transcription (43), there is the cooperation of a 5 nanotube super-aggregates (Figure 1). This super-super aggregation is similar to a recently produced semiconducting material, having an arrangement made by π−π stacking aromatic moieties supported by bordering H bonds that, in the whole, constitute flanked columnar structures (44).

The elemental components of a carrier device were imagined to compose as a cyclic structure formed by the super-assembly of 8 phosphate groups linked by H bonds (Figure 5). That was in accordance with what reported by Hossain et al. who characterized a cyclic octamer of dihydrogen phosphates by electro spray ionization mass spectrometry and 31 P nuclear magnetic resonance (28). These modular aggregates have the assets of an agile structure that can self-assemble by hooking at the amino-groups of the protein that has to be carried. The cooperation of several modules in protein carrying has to be considered mandatory since, in Z-DNA form, the involvement of 5 flanked nanotubes has been demonstrated (6-8, 10, 11,14,15).

As stated before, a negative electric arrangement of the protein interacting side is indispensable for the activation of the motive apparatus. HMGB1 proteins, histones and, in general, all binding proteins have two major possibilities for modifying the charge of their interactive sites by means of phosphate groups: to electrostatically interact (negatively - 2 e- - charged phosphate ions vs. positively - 1 H+ - charged amino groups) and/or to become phosphorylated in the amino acid serine, lysine, and threonine (45-48). Both these options are able to activate the establishing of H bonds among phosphate-phosphate groups. In fact, a protein-linked phosphate group maintains its propensity of establishing H bonds with the reactive groups within reach (49-51).

Plainly, the system could function as an electric motored bogie that carries the protein along the DNA up to a befitting destination, the sequence-specific DNA-binding domain (52) by means of a structure made of phosphates, so configuring a sort of robotic handling of the task.

#### Rotational motion

This kind of linear motion has, potentially, consequences in the rotation of histones too, since the angular momentum is the rotational equivalent of the linear one, and the motion of a “body” under a central force always remains in the plane defined by its initial position and velocity (53).

The nucleosome is formed by histones H2A, H2B, H3, and H4, with each contributing two molecules, so forming a histone octamer around which DNA wraps in approximate two turns in a left-handed way. H1 (H5) is a linker histone, which binds at the point where DNA enters and exits the nucleosome. Positive charges, differently from negative ones, are determinant in contrasting the histone rotation as evinced by the fact that the deletion of positive charges in histone tails facilitated partial DNA unwrapping, most significantly for H3 and H2B (54).

The structure has theoretically rotating abilities in the anti-clockwise direction that permit the histone octamer to slide towards the DNA loop (55,56). However, as evinced by an exhaustive recent review, the histone dynamics is currently considered a puzzling matter (57). Briefly, Authors affirm that, within the current knowledge, a “sliding” motion of the DNA around the octamer, is far too expensive as it would require that all binding sites of the strands break at the same time and the “rolling” motion of the octamer, obtained breaking sites at one end and closing sites on the other, is also not possible because a fully wrapped nucleosome has no sites to roll on.

The acknowledgment of the NAP carrying structure would shed light on this topic: the same system hypothesized for the protein linear motion could be advocated for the rotation of histones without the necessity of supposing any break on DNA strands. In other words, the NAP - DNA loop would function as a coil around a proteic core in which the linear motion of phosphates, exalted by the increased electromagnetic field (58) produces the octamer rotation and, consequently activates the wrapping and rewrapping dynamics of the DNA strands.

## CONCLUSION

The theoretical parts of this paper have been constructed following the principle that, if something exists (a nanotube system structured as a n-type semiconductor enveloping the double strands) has to serve to something that, in consideration of its dimension and structural features, cannot be of scarce bio-physical relevance. At light of the NAP acknowledged influence, many incongruences of the current bio-physical interpretations of the motion of protein and histones could be more directly explained.

When wrapping the DNA, NAPs accomplish structural and functional missions in cooperation with the double strands and have to be, therefore, considered a not secondary example of natural self-organizing biostructures. Their assembly, based on the supramolecular principles, allows that they function as smart polymers having electronic abilities.

The functioning of NAPs as semiconductors has to be still explored, but, since the bio-physical presuppositions here described are, in my opinion, well represented and "electrifying", it is easy to foresee that a new wide scenario will open to the scientists that will be engaged in the study. Furthermore, this functional supramolecular polymeric aggregation has the assets for becoming a template for the production of electronic nano-constructed materials.

In a wider prospect, it is possible to affirm that time has arrived for reconsidering the biochemical processes at the light of the phosphate-phosphate interaction, a novelty that promises to change our traditional beliefs on many aspects of cell biology.

## ACKNOWLEDGEMENTS

This paper is dedicated to the people that worked with me.

## ABBREVIATIONS

NAP: nuclear aggregate of polyamine
EF: electric field
H bond: hydrogen bond
MALDI-TOF MS: Matrix-assisted laser desorption ionization time-of-flight mass spectrometry
AFM: atomic force microscopy
HMGB1: high mobility group box 1
H bond: hydrogen bond

## REFERENCES

1. Xie, L-H, Q-D. Ling, X-Y Hou, W. Huang. 2008. An Effective Friedel–Crafts Postfunctionalization of Poly(N-vinylcarbazole) to Tune Carrier Transportation of Supramolecular Organic Semiconductors Based on π-Stacked Polymers for Nonvolatile Flash Memory Cell. J. Am. Chem. Soc. 130: 2120–2121.

2. Berlin, Y. A., A. A. Voityuk, and M. A. Ratner. 2012. DNA Base Pair Stacks with High Electric Conductance: A Systematic Structural Search. ACS Nano. 6: 8216–8225.

3. Chaoren, L., C. Liu, L. Xiang, Y. Zhang, P. Zhang, D.N. Beratan, Y. Li, N. Tao. 2016. Engineering nanometre-scale coherence in soft matter. Nature Chemistry 8: 941–945.

4. Genereux, J.C. and J. K. Barton. 2010. Mechanisms for DNA Charge Transport. Chem. Rev. 110:1642–1662.

5. Hong, F., F. Zhang, Y. Liu, H. Yan. 2017. DNA Origami: Scaffolds for Creating Higher Order Structures. Chem. Rev. 117:12584–12640.

6. D’Agostino, L. and A. Di Luccia. 2002. Polyamines interact with DNA as molecular aggregates. Eur. J. Biochem./FEBS. 269:4317–4325.

7. D’Agostino, L., M. di Pietro, and A. Di Luccia. 2005. Nuclear aggregates of polyamines are supramolecular structures that play a crucial role in genomic DNA protection and conformation. FEBS J. 272: 3777–3787.

8. D’Agostino, L.,M. di Pietro, and A. Di Luccia. 2006. Nuclear aggregates of polyamines., IUBMB Life. 58:75–82.

9. Di Luccia, A., G. Picariello, G. Iacomino, A. Formisano, L. Paduano, L. D’Agostino. 2009. The in vitro nuclear aggregates of polyamines. FEBS J. 276: 2324–2335.

10. Iacomino, G., G. Picariello, F. Sbrana, A. Di Luccia, R. Raiteri, L. D’Agostino. 2011. DNA is wrapped by the nuclear aggregates of polyamines: the imaging evidence. Biomacromolecules 12:1178–1186.

11. Iacomino, G., G. Picariello, and L. D’Agostino. 2012. DNA and nuclear aggregates of polyamines, Biochim. Biophys. Acta 1823:1745–1755.

12. Iacomino G., G. Picariello, I. Stillitano, L. D’Agostino. 2014. Nuclear aggregates of polyamines in a radiation-induced DNA damage model, Int. J. Biochem. Cell Biol. 47: 11–19.

13. Picariello, G., G. Iacomino, A. Di Luccia, L. D’Agostino 2014. Mass spectrometric analysis of in vitro nuclear aggregates of polyamines, Rapid Commun. Mass Spectrom. 28: 499–504.

14. Picariello, G., G. Iacomino, and L. D’Agostino. 2015. Phosphate-induced polyamine self-assembly. In Encyclopedia of Biomedical Polymers and Polymeric Biomaterials. M. Mishra, editor. Taylor & Francis, New York, pp. 5951–5964.

15. Iacomino G., G. Picariello, F. Sbrana, R. Raiteri, L. D’Agostino. 2016. DNA-HMGB1 interaction: The nuclear aggregates of polyamine mediation. Biochim. Biophys. Acta - Proteins and Proteomics 1864: 1402–1410.

16. Lomozik, L., A. Gasowska, R. Bregier-Jarzebowska, R. Jastrzab. 2005. Coordination chemistry of polyamines and their interactions in ternary systems including metal ions, nucleosides and nucleotides. Coord. Chem. Rev. 249: 2335–2350.

17. Lutz, K., C. Groger, M. Sumper, E. Brunner. 2005. Biomimetic silica formation: Analysis of the phosphate-induced self-assembly of polyamines. Phys. Chem. Chem. Phys. 7: 2812–2815.

18. Laucirica, G., W. A. Marmisollé, and O. Azzaroni. 2017. Dangerous liaisons: anion-induced protonation in phosphate–polyamine interactions and their implications for the charge states of biologically relevant surfaces. Phys. Chem. Chem. Phys. 19: 8612–8620.

19. Fenniri, H., P. Mathivanan, K. L. Vidale, D. M. Sherman, K., Hallenga, K. V. Wood, J. G. Stowell. 2001. Helical Rosette Nanotubes: Design, Self-Assembly, and Characterization. J. Am. Chem. Soc. 123:3854–3855.

20. Keizer, H. M., and R. P. Sijbesma. 2005. Hierarchical self-assembly of columnar aggregates. Chem. Soc. Rev. 34: 226–234.

21. Bhalla, V., R. P. Bajpai, and L. M. Bharadwaj. 2003. DNA electronics. EMBO reports 4: 442–445.

22. Inokuchi, H. 2006. The discovery of organic semiconductors. Its light and shadow. Organic Electronics 7:62–76.

23. Kenichi, F. 1981. The Role of Frontier Orbitals in Chemical Reactions. Nobel lecture.

24. Rios, C. and R. Salcedo. 2014. Molecules 19:3274–3296.

25. Jeppesen, J.O., M. Brøndsted Nielsen, and J. Becher. 2004. Tetrathiafulvalene cyclophanes, and cage molecules. Chem. Rev. 104: 5115–5131.

26. Singh, A. K. 2016. Physicochemical, Electronic, and Mechanical Properties of Nanoparticles. In Engineered Nanoparticles. A. K. Singh, editor. Elsevier Academic Press, San Diego, pp. 77–123.

27. Mima, T., T. Kinjo, S. Yamakawa, R. Asahi. 2017. Study of the conformation of polyelectrolyte aggregates using coarse-grained molecular dynamics simulation. Soft Matter 13:5991–5999.

28. Hossain, M.A., M. Isiklan, A. Pramanik, M.A. Saeed, F.R Fronczek,. 2012. Anion cluster: Assembly of dihydrogen phosphates for the formation of a cyclic anion octamer. Cryst. Growth Des. 12:567–571.

29. Mata, I., I. Alkorta, E. Molins, E. Espinosa. 2012. Electrostatics at the origin of the stability of phosphate-phosphate complexes locked by hydrogen bonds. Chemphyschem, 13:1421–1424.

30. Ross, E.D., P.R. Hardwidge, and L.J.3rd Maher. 2001. HMG proteins and DNA flexibility in transcription activation, Mol. Cell. Biol. 21: 6598–6605.

31. Mata, I., I. Alkorta, E. Molins, E. Espinosa, 2013. Tracing environment effects that influence the stability of anion-anion complexes: The case of phosphate-phosphate interactions. Chem. Phys. Lett. 555: 106–109.

32. Mata, I., E. Molins, I. Alkorta, E. Espinosa. 2015. The paradox of hydrogen-bonded anion-anion aggregates in oxoanions: a fundamental electrostatic problem explained in terms of electrophilic…nucleophilic interactions. J. Phys. Chem. A 119: 183–194.

33. Tyner, K.M., R. Kopelman, and M.A. Philbert. 2007. “Nanosized Voltmeter” Enables Cellular-Wide Electric Field Mapping. Biophysical J. 93: 1163–1174.

34. Cunningham, J., V. Estrella, M. Lloyd, R. Gillies, B. R. Frieden, R Gatenby. 2012. Intracellular Electric Field and pH Optimize Protein Localization and Movement. PLoS ONE 7: e36894.

35. Giorgio, V., S. von Stockum, M. Antoniel, A. Fabbro, F. Fogolari, M. Forte, G. D. Glick, V. Petronilli, M. Zoratti, I. Szabó, G. Lippe, P. Bernardi. 2013. Dimers of mitochondrial ATP synthase form the permeability transition pore. PNAS 110: 5887–5892.

36. Halestrap, A. P. and A. P. Richardson. 2015. The mitochondrial permeability transition: A current perspective on its identity and role in ischemia/reperfusion injury. J. Mol. Cell. Cardiol. 78:129–141.

37. O’Reilly, C.M., K.E. Fogarty, R.M. Drummond, R.A. Tuft, J.V. Walsh. 2003. Quantitative Analysis of Spontaneous Mitochondrial Depolarizations. Biophysical J. 85:3350–3357.

38. Loewenstein, W. R. and Y. Kanno. 1962. Some electrical properties of the membrane of a cell nucleus. Nature 195:462–464.

39. Mazzanti, M., L. J. DeFelice, J. Cohen, H. Malter. 1990. Ion channels in the nuclear envelope. Nature 343:764–767.

40. Oberleithner, H., S. Wünsch, and S. Schneider. 1992. Patchy accumulation of apical Na+ transporters allows cross-talk between extracellular space and cell nucleus. PNAS 89:241–245.

41. Mazzanti, M., J.O. Bustamante, and H. Oberleithner. 2001. Electrical Dimension of the Nuclear Envelope. Physiol. Rev. 81:1–19.

42. Lin, X., M. Suzuki, M. Gushiken, M. Yamauchi, T. Karatsu, T. Kizaki, Y. Tani, K. Nakayama, M. Suzuki, H. Yamada, T. Kajitani, T. Fukushima, Y. Kikkawa, S. Yagai. 2017. High-fidelity self-assembly pathways for hydrogen-bonding molecular semiconductors. Scientific Reports 7:43098.

43. Rich, A., and S. Zhang. 2003. Timeline: Z-DNA: the long road to biological function. Nat. Rev. Genet. 4:566–572.

44. Gorbunov, A. V., A. T. Haedler, T. Putzeys, R. H. Zha, H-W. Schmidt, M. Kivala, I. Urbanavičiūtė, M. Wübbenhorst, E. W. Meijer. and M. Kemerink. 2016. Switchable Charge Injection Barrier in an Organic Supramolecular Semiconductor. ACS Appl. Mater. Interfaces 8:15535–15542.

45. Youn, J. H. and J.-S. Shin. 2006. Nucleocytoplasmic Shuttling of HMGB1 Is Regulated by Phosphorylation That Redirects It toward Secretion. J. Immunol. 177:7889–7897.

46. Oh, Y. J., J. H. Youn, Y. Ji, S. E. Lee, K. J. Lim, J.E. Choi, J.-S Shin,. 2009. HMGB1 Is Phosphorylated by Classical Protein Kinase C and Is Secreted by a Calcium-Dependent Mechanism. J. Immunol. 182:5800–5809.

47. Bannister, A. J., and T. Kouzarides. 2011. Regulation of chromatin by histone modifications. Cell Research 21:381–395.

48. Rossetto, D., N. Avvakumov, and J. Côté. 2012. Histone phosphorylation. A chromatin modification involved in diverse nuclear events. Epigenetics 7: 1098–1108.

49. Mandell, D.J., I. Chorny, E. S. Groban, S. E. Wong, E. Levine, C. S. Rapp, M. P. Jacobson. 2007. Strengths of Hydrogen Bonds Involving Phosphorylated Amino Acid Side Chains. J. Am. Chem. Soc. 129: 820–827.

50. Hunter, T. 2012. Why nature chose phosphate to modify proteins. Phil. Trans. R. Soc. B 367:2513–2516.

51. Thapar, R. 2014. Contribution of protein phosphorylation to the binding-induced folding of the SLBP–histone mRNA complex probed by phosphorus-31 NMR. FEBS Open Bio 4: 853–857.

52. Krajewska, W. M. 1992. Regulation of transcription in eukaryotes by DNA-binding proteins Int. J. Biochem. 24:1885–1898.

53. Goldstein, H. 1980. Classical Mechanics, 2nd ed., Addison-Wesley, Reading.

54. Kenzaki, H., and S. Takada. 2015. Partial Unwrapping and Histone Tail Dynamics in Nucleosome Revealed by Coarse-Grained Molecular Simulations. PLOS Computational Biology DOI:10.1371/journal.pcbi.1004443.

55. Li, W., S.X. Dou, and P.Y. Wang. 2005. The histone octamer influences the wrapping direction of DNA on it: Brownian dynamics simulation of the nucleosome chirality. J. Theor. Biol. 235:365–372.

56. Li. W., S. X. Dou, P. Xie, P. Y. Wang. 2006. Brownian dynamics simulation of directional sliding of histone octamers caused by DNA bending. Phys. Rev. E 73:051909.

57. Eslami-Mossallam, B., H. Schiessel, and J. van Noort. 2016. Nucleosome dynamics: Sequence matters. Adv.Colloid Interface Sci. 232: 101–113.

58. Livingston, J. D. 1996. Driving Force, The Natural Magic of Magnets, first ed., Harvard University Press, Cambridge.

